# Simulation of population development in the predator-prey system under changing conditions of interaction

**DOI:** 10.1101/016691

**Authors:** Alexandr N. Tetearing

## Abstract

The numerical models of populations behaviour, simulated under changing condition of populations interaction (the hungry or full-fed populations, the existence of persons in the personal areas or in the common territory), demonstrate the increase, reduction or damping of oscillations amplitude, that corresponds qualitatively to known experimental data, obtained for real population of bacteria.

---

The predator-prey equations systems can be successfully used for description and simulation of the real natural biological systems.

The process of simulation of these systems is rather complicated by reason of the presence in system of a large number of parameters and coefficients, and by reason of periodic interleaving of the several existence modes of populations interacting, through which these predator-prey systems pass in their development. The simulation of the population interaction can be well executed with using the programme numerical methods, applicable for solution of the differential equations systems.

As example of this simulation we will consider the simplest case of the description of the species interaction process in the predator-prey population system.

We simulate a system that describes, for example, the population development of bacteria – the protozoa unicellular organisms, that reproduce by cell fission.

The most popular example that is found practically in every biology article on this subject, related to the predator-prey population interaction, is the system of the differential equations by Lotka-Volterra:

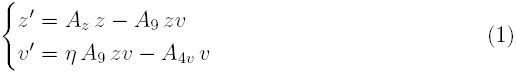

Here, the time-depended functions *z* and *v* are, respectively, the total population mass of the prey and predators. The *η*-coefficient is always positive and less than one.

We use also the following designations:

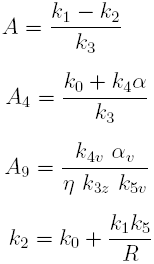

The addition of the badge *z* to a some coefficient denotes, that this coefficient pertains to population of prey (the coefficients, characterizing the populations of prey and predators, have different values).

*R* is the amount of food resource, provided by environment per unit time on the one unit of the population habitat area.

All the *k*_*i*_ are the constant factors.

This system of equations describes a certain stage of development of predatorprey system at interaction of two populations of unicellular organisms (the bacteria).

The classical graph of these two periodic functions is shown in figure 1.

**Figure 1:**
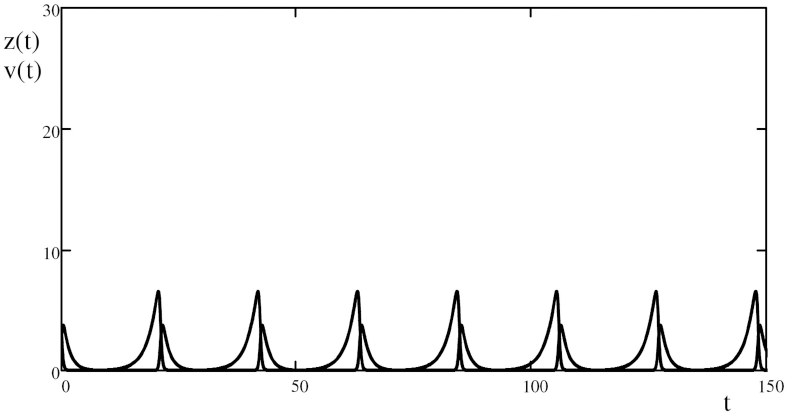
Curves of population mass change for predator population *v*(*t*) and prey population *z*(*t*), according to equations system 1.

The solutions of the Lotka-Volterra equations system (1) are normally depicted in this form.

For plotting of these functions graphs we used the following parameters values:

*k*_0*z*_ = 0.1, *k*_1*z*_ = 0.3, *k*_2*z*_ = 0.16, *k*_3*z*_ = 0.3, *k*_4*z*_ = 0.3, *k*_5*z*_ = 0.2, *k*_6*z*_ = 0.02, *α*_*z*_ = 0.8, *A*_*z*_ = 0.467, *A*_1*z*_ = 0.667, *A*_2*z*_ = 0.02, *R*_*z*_ = 1, *k*_0*v*_ = 0.1, *k*_1*v*_ = 1, *k*_3*v*_ = 0.3, *k*_4*v*_ = 0.3, *k*_5*v*_ = 0.5, *k*_6*v*_ = 0.03, *α*_*v*_ = 0.4, *A*_1*v*_ = 3, *η* = 0.65, *K*_1_ = 5.128, *A*_7_ = 1.667, *A*_9_ = 1.231.

The factors with index *z* pertain to the prey population, the factors with index *v* pertain to predators population. The coefficient redundancy is necessary for graph drawing of the population development functions under conditions of the successive modes of their existence (in particular, for checking of the transition mode of population development); besides, the coefficient redundancy is useful for independent reproduction of these numerical graphs by our readers.

The graph of function of the prey population mass changes *z*(*t*), marked in figure as “Prey”, has a more high amplitude of the oscillations, and graph of function of the predator population mass changes *v*(*t*), marked in figure by “Predator” inscription, has smaller value of the maximum, and furthermore, there is a time lag between the function *z*(*t*) maximum and the function *v*(*t*) maximum.

The figure 2, on a large scale, shows the one period of *z*(*t*) and *v*(*t*) functions to see better the typical form of these curves.

**Figure 2:**
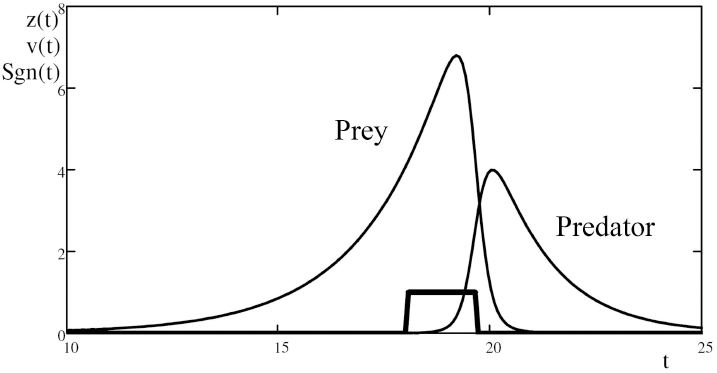
The predator *v*(*t*) and prey *z*(*t*) mass population changes, corresponding to the equations system 1, presented in more large scale. The square-wave stair is the signal function *Sgn*(*t*) (it is shown by a thick line).

The initial values of the population mass for prey *z*_0_ and predator *v*_0_ are: *z*_0_ = 3 and *v*_0_ = 3.

The curves are built with software, with use the fourth order Runge-Kutta method with a fixed step.

We assume, that the Lotka-Volterra equation system (1) describes the population interaction of representatives of full-fed prey population, that live in the personal territory areas, and hungry predators, that also live in their personal areas.

The person existence in a individual area means that the person has his own territory for hunt and feeding, and the area is not crossed with other persons hunt areas.

The hungry existence of persons means that persons consume the food less, than could eat under more happy circumstances (at food profusion), and have to lead for all time the permanent hunt for a food resource.

But this is because, the number of persons in the population is changing, practically always a such moment will come, when the number of predators becomes small in compare with number of prey, and predator population goes (for a short time) into full-fed existence mode.

Herewith, the Lotka-Volterra equation system (1) already does not work, and we need to use for description of populations interaction the equation system, that describes the full-fed prey and full-fed predators in their personal areas:

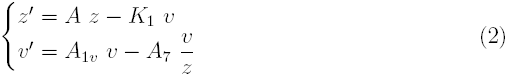

We use here the following designations:

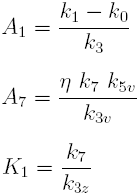

How do we define the moment of time, when, under serial calculations, we need to replace the first equation system to the another?

For this, when the predator population mass is changing, we need to check every time for predator population the conditions of the transition from hungry existence mode to the full-fed existence mode.

The condition of transition from the hungry existence mode to the full-fed mode for the predator population is an inequality:

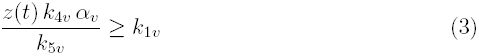

Using the condition (3) we define the signal function *Sgn*(*t*), which is equal to one, if (3) condition is satisfied (the full-fed predators); and it is equal to zero – if the condition is not satisfied (the predators are hungry).

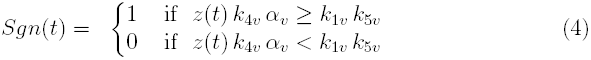

This signal function will mark on the time axis the time intervals, when the predators live a full-fed existence.

These full-fed intervals of a predator existence occur quite frequently – this happens whenever the number of predators is sufficiently small, and the number of prey is sufficiently large.

The graph of the *Sgn*(*t*) function is shown in figure 2 as square-wave stair with unit-equaled height, drawn with the thick line. On this marked time interval, at calculations of function *z*(*t*) and *v*(*t*) values, we need to use the (2) equation system instead of (1).

In spite of the fact that these intervals are continued for a short time, they make the essential changes to *z*(*t*) and *v*(*t*) functions. With taking into account these short-time full-fed modes of predators existence, here the smooth periodic growth of the prey population mass comes at time of each following periodic maximum. These changes in the *z*(*t*) and *v*(*t*) functions are shown in figure 3.

**Figure 3:**
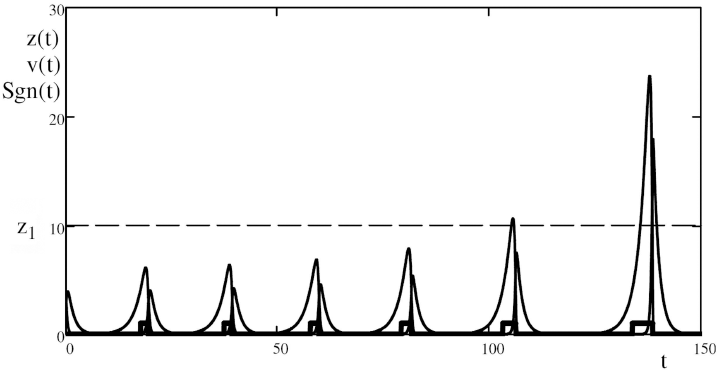
Mass change curves for the predators *v*(*t*) and prey *z*(*t*) populations constructed with taking into account the presence of the full-fed mode periods in the predators population.

The graphs in figures 1 and 3 are given in the same scale, in order to better see the changes, associated with the presence the full-fed existence periods in predators population.

The graph 3 shows, how the amplitude of periodic oscillations in number of prey and predators increases with time. The period of the oscillations of population mass is also increasing for both populations.

Besides, the graph shows that with time, from one period to the other, the duration of full-fed period for the predator population also elongates (these periods are shown in figure as square-wave stairs with unit-equaled height, marked by a thick line).

Under these conditions, without any external factors limiting population growth, both populations will grow unlimitedly with time.

In practice, these are the factors, limiting population growth, such as the lack of food resources or the territorial limitation of population habitat area.

So far as the experiences on study of the bacteria populations interaction are usually conducted in the laboratory, for the further illumination of our problem we shall choose, as the limiting factor, the finite sizes of laboratory dishes.

By reason of increase in number of bacteria in the laboratory dish (we shall consider it as the flat container with nutrient medium layer with one-bacterium thickness) the increase of bacteria density occurs in the test-dish, and the bacterias (which were feeding in the personal areas before) are now beginning to compete with each other for food resource in the common territory.

We assume (in our case) that the prey population will face with these territorial restrictions earlier than the predators population.

When the population of prey passed from the conditions of the full-fed existence in the personal areas to the conditions of the full-fed existence in the common territory, the equation system in our problem, describing interaction of populations, should be changed.

The predator-prey population interaction, in this case, will be described by the new equations.

If predators are hungry, we have to use the following equations:

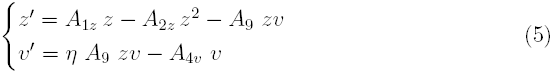

If the predators are full-fed, we have to use the following equations:

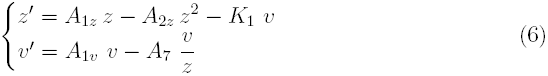

Here *A*_2_ is:

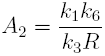

The condition for the prey population transition from full-fed mode with existence in the personal areas to the full-fed mode with existence in the common territory is the inequality:

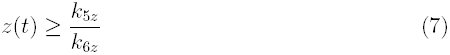

For our case:

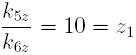

That is to say, as soon as the graph of *z*(*t*) function rises above the horizontal dashed line *z* = 10 in figure 3, the prey population goes to the mode of existence in the common territory.

The graphs of *z*(*t*) and *v*(*t*) functions, that are drawn taking into account this limiting factor for population of the prey, are shown in figure 4.

**Figure 4:**
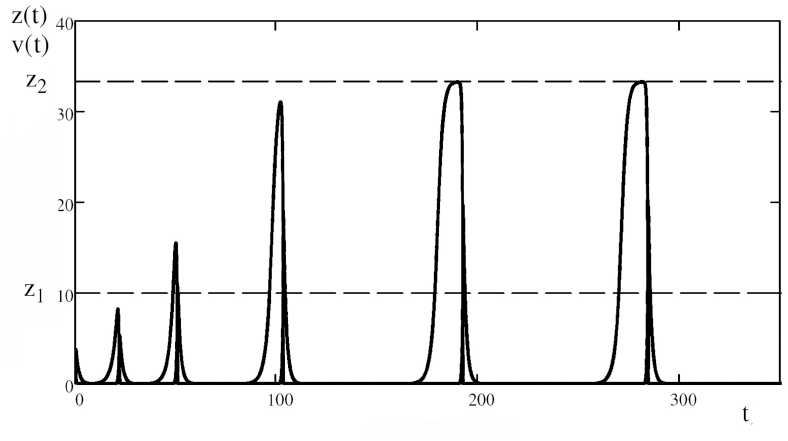
Graph of the predators *v*(*t*) and prey *z*(*t*) population mass changes, that are drawn taking into account the territory limitation factor for population of the prey.

The graph shows, that after the prey population reaches of a certain limiting value *z*_2_ in its maximum, the maximal value of the prey population mass in the following oscillations doesn’t change with time. Also, from this time point, the period of oscillation of the both populations mass stabilizes and becomes constant.

The value of the limit *z*_2_ can be determined using the formula:

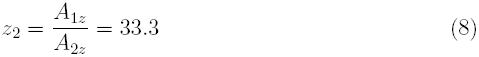

Comparing graph 1 and 2 with graphs 4 and 5 (that are corrected with taking into account the territory limitation factor for population of the prey) we see that, with our adjustments, the function graphs have changed greatly. Namely, the period and amplitude of the both populations mass oscillations changed.

In figure (5) the prey population graph (the function *z*(*t*)) has the flat plateau instead of sharp top-maximum (on this plateau the graph of function keeps a some time a certain constant limit value *z*_2_). The width of this plateau (time duration) is defined by the initial parameters of the both populations development and (for some parameters) it can last a very long time.

**Figure 5:**
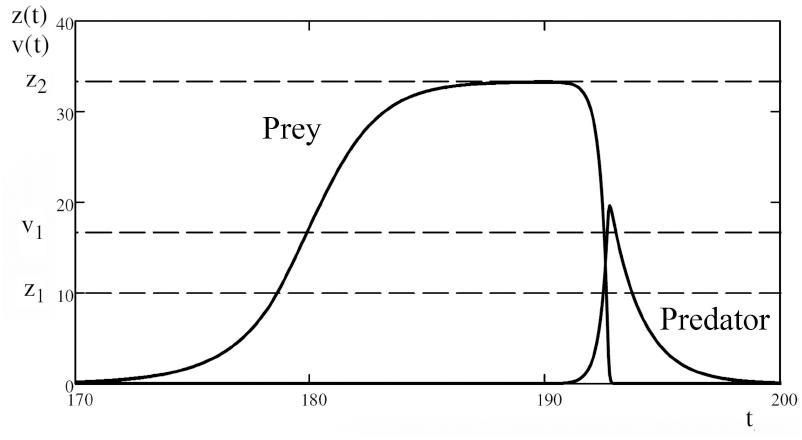
Graph of the predators *v*(*t*) and prey *z*(*t*) population mass changes, that are drawn taking into account the territory limitation factor for population of the prey (in a large scale).

However, in spite of the significant changes, which have taken place in our population development functions in this article, this is yet not the complete set of parameters, which we need to take into account at description of the similar interaction of the two populations.

In particular, we expect that persons in population of the prey always live a full-fed existence (sometimes despite the overpopulation, which leads to the increase of population energy expenses on the hunting for food resources). Consequently, we need to check constantly the full-fed existence condition for the prey population.

In our case, this condition is:

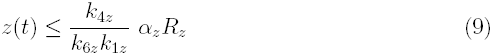

We shall substitute in left-hand side of the inequality the maximal mass value *z*_2_, which population of the prey reaches in its development, and shall calculate the right-hand side of the inequality:

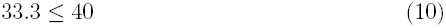

It is seen that this condition in our case is always satisfied (the prey in our figure lives always a full-fed existence).

If this condition is violated periodically, we would change the our equations under calculations at this time (to change these equations in accordance with conditions for hungry development of prey population in the common territory).

In this article we have chosen the parameters values deliberately by such way, to show the simplest example for the solution of problem on the predator-prey population interaction.

Now is the time to remember the predators population. The predators, when the number of individuals in population is increased, also can face the factor of restriction of a population growth. This factor, in our case, will also cause the periodic territorial overpopulation of predators. At times (with a large number of predators) the predators population will pass from initial mode of existence in the personal habitat areas to the mode of existence in the common territory.

Herewith, the equations, that describe the interaction of the predator-prey populations system, will changes appropriately.

The condition of predators population transition from the hungry mode of existence in the personal areas to the hungry mode of existence in the common territory is the condition:

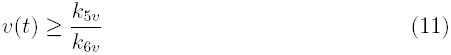

In our case:

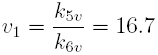

The *v*_1_ value is marked in figure 5 by a horizontal dashed line (the middle line of the three lines shown in figure)

The figure shows that graph of function *v*(*t*) (the function with “Predator” mark) sometimes rises above horizontal dashed line *v* = *v*_1_ that corresponds to transition of predators population into the mode of existence in the common territory. For this time interval we need again to change our current system of the equations.

If the prey inhabit in the common territory, we should use the following system of equations:

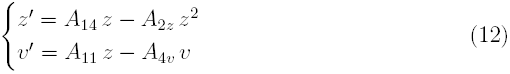

If it is the time, when the prey dwell in the personal areas, we use the equations:

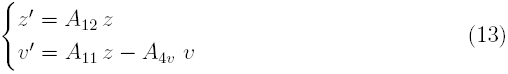

We use here the following designations:

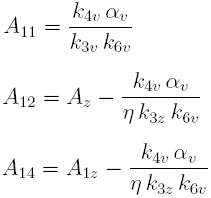

And, certainly, as soon as the predators population mass will fall below value *v* = *v*_1_, we should return again to the equations, which we used before predators transition into the mode of existence in the common territory.

The final graphs of the functions, describing the system of the predator-prey populations, are presented in figures 6 and 7 (these are the same graphs, but on a larger scale).

**Figure 6:**
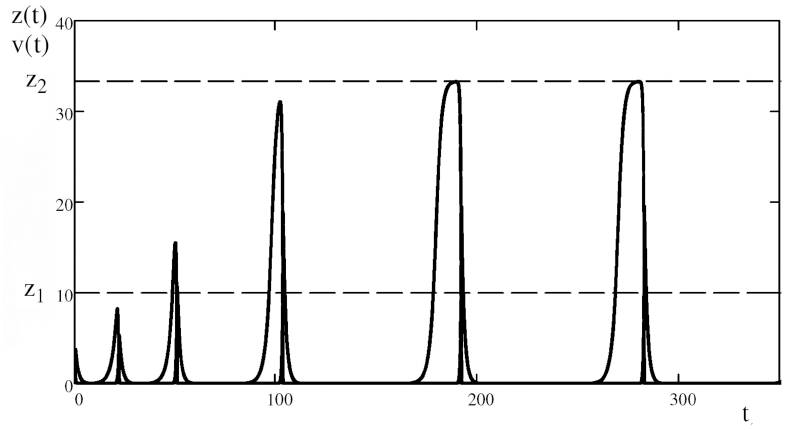
Graphs of the population mass change for predators *v*(*t*) and prey *z*(*t*), that are builded taking into account the territory limitation for the predators population.

**Figure 7:**
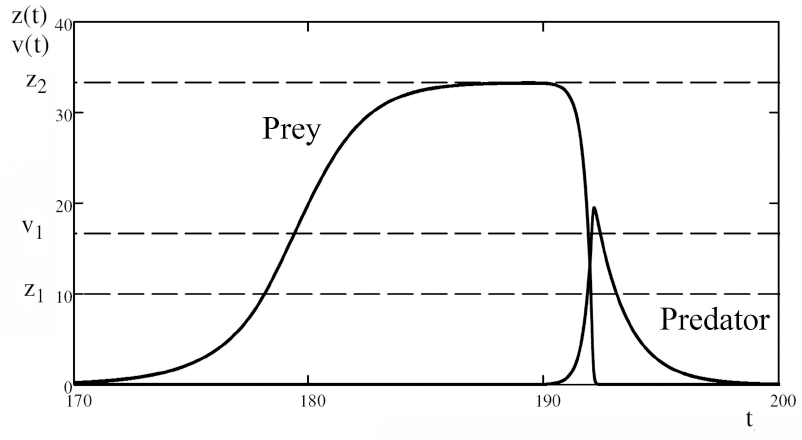
Graphs of the population mass change for predators *v*(*t*) and prey *z*(*t*), that are builded taking into account the territory limitation for the predators population (on a larger scale).

Since, in this case, the predators population exists in the mode of existence in the common territory during the periodic short-time intervals, the changes in the graphs are not significant. However, when comparing figures 4 and 5 with figures 6 and 7, we can see that the periodic maximums of the functions of changes in the population mass for the prey and the predators, are shifted to the left on axis of time. Besides, the period oscillations of the both populations mass has changed little (the period decreased).

This can be explained by the fact, that in the common territory the persons of predators population spend more time and energy on the hunt for food resources. For this reason the overall mass increment of the predators population is reduced. The prey population, opposite, under these conditions, restores faster the maximum value of its size because the number of predators has decreased.

Now we will give another example of the populations interaction for predatorprey systems.

In this case we will change the experiment conditions, that are described by our mathematical model.

Assume that at the initial time the prey population exists under conditions of the food resource profusion and with the full absence of predators. In this case, as we know, the prey population reaches its maximal number of individuals and hereinafter (with the absence of predators) the number of persons in the prey population does not change.

After the population reaches its maximum, we put into laboratory dish with prey a small number of predators, and at this point of time we begin to build the mathematical graphs, that describe the populations interaction.

Unlike previous example of the populations development, in this case the predators fall immediately into conditions of full-fed existence on the personal areas and the population interaction should be described initially by the following system of equations:

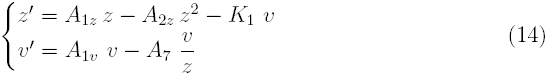

This equations system describes the full-fed prey in the common territory and the full-fed predators in the personal areas.

Hereinafter, at this predator-prey system simulation, as before, we will change the current equations system in accordance to problem of the base classification, whenever the conditions of the existence of predators or prey change.

At predator-prey system setting we use the following values of parameters and coefficients:

*k*_0*z*_ = 0.1, *k*_1*z*_ = 0.3, *k*_2*z*_ = 0.16, *k*_3*z*_ = 0.3, *k*_4*z*_ = 0.3, *k*_5*z*_ = 0.2, *k*_6*z*_ = 0.02, *α*_*z*_ = 0.8, *A*_*z*_ = 0.467, *A*_1*z*_ = 0.667, *A*_2*z*_ = 0.02, *R*_*z*_ = 1, *k*_0*v*_ = 0.1, *k*_1*v*_ = 1, *k*_3*v*_ = 0.3, *k*_4*v*_ = 0.3, *k*_5*v*_ = 0.75, *k*_6*v*_ = 0.1, *α*_*v*_ = 0.2, *A*_1*v*_ = 3, η = 0.4, *K*_1_ = 8.333, *A*_7_ = 2.5, *A*_9_ = 0.667.

The factors with index *z* pertain to the prey population, the factors with index *v* pertain to the predators populations.

The initial mass *z*_0_ of the prey population (the full-fed mode of population existence in the common territory) in this experiment is defined by value *z*_2_, which is equal to:

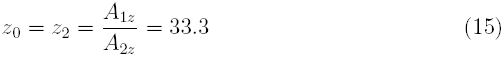

The initial mass of the predators population (the full-fed predators in the personal areas) is *v*_0_ = 0.5

The graphs, which illustrate the populations interaction under these parameters values, are shown in figure 8.

**Figure 8:**
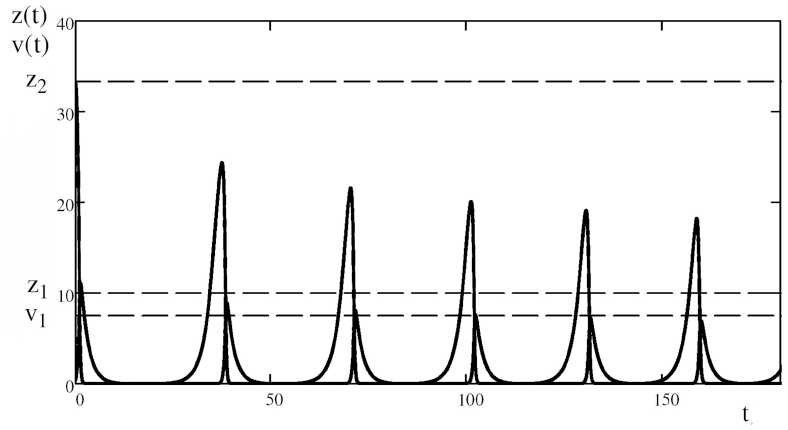
Graphs of the populations interaction for predators *v*(*t*) and prey *z*(*t*) that are correspond to the equations system 14.

Figure 9 shows the same graphs on a larger scale. It shows the first period of the populations mass oscillations, that is started at time *t* = 0.

**Figure 9:**
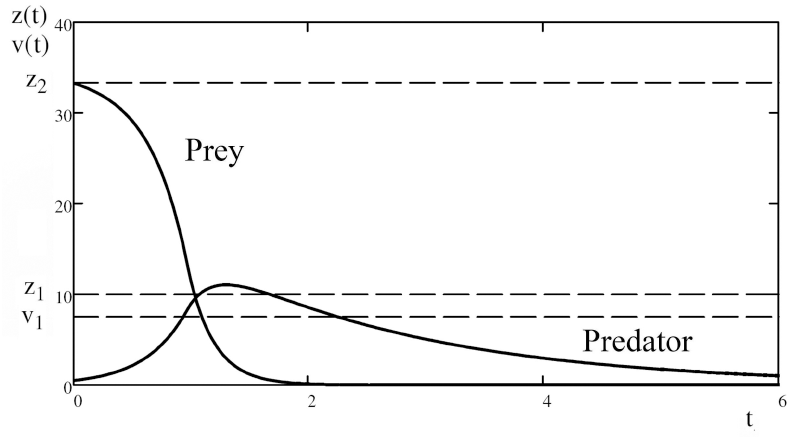
Graphs of the populations interaction for predators *v*(*t*) and prey *z*(*t*) that are correspond to the equations system 14 (on a larger scale). Part of a first period of the populations mass oscillations.

Let’s try to compare the obtained graphs with the practical results of the laboratory studies.

The experiences on the bacteria populations development in the predator-prey system were carried out for the first time by Georgy Frants Gause (the doctor of the biological sciences, graduated from the biological branch of physics and mathematics faculty of the Moscow university).

The results of experience were published by scientist in his book “The Struggle for Existence” [3], published in 1934. At that time, the equation system (1), that was offered by Lotka and Volterra, has been well known in biology, and Gause conducted his laboratory experiments in order to verify in practice these mathematical equations.

The scientist experimented with bacteria populations and planned to fix in laboratory experience the cyclical oscillations in the number of predator and prey, that were prescribed by solutions of the Lotka-Volterra equation system (1).

The data, obtained as a result of experiment and published in the book [3], are shown on figure 10.

**Figure 10:**
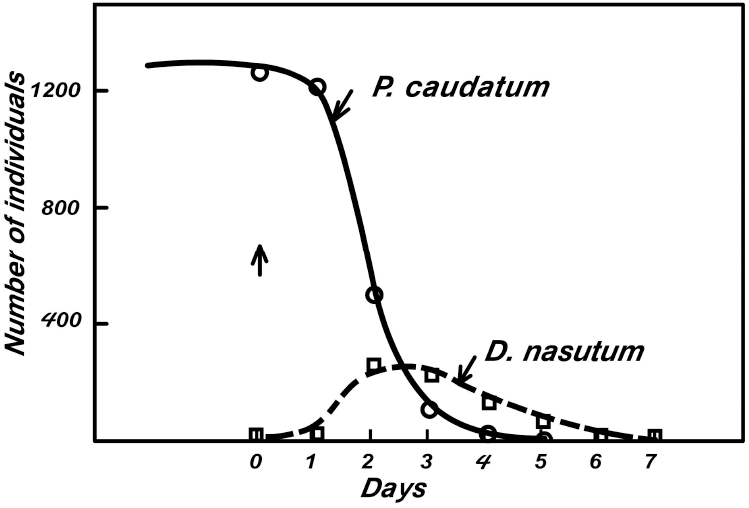
Graphs of interaction of predators (the Didinium nasutum bacteria) and prey (the Paramecium caudatum bacteria). The illustration taken from the book “The Struggle for Existence” by F.G.Gause [3].

The vertical axis represents the number (in individuals) of predators and prey, the horizontal axis represents the time (in days).

As the predator-prey model, Gause considered the interaction of the bacteria populations: the Didinium nasutum (as the predators) and the Paramecium caudatum (as the prey).

As a food for prey population was used the saline medium of Osterhout with the Bacillus pyocyaneus. The feeding medium was given in equal amounts and was replaced daily, with removing of the products of vital activity of microorganisms – to provide for the prey population (Paramecium caudatum) the sufficient nutrition and suitable conditions for intensive reproduction.

The predators population ate the representatives of the Paramecium caudatum.

The results of the several experiences demonstrated that prey population is always destroyed completely by predators, then, after awhile, the predators population dies too. Thereby Gause concluded, that the Lotka-Volterra’s mathematical model of the populations interaction is invalid (it is inapplicable in this case), and the periodic oscillations in the number of individuals in both populations don’t occur here.

The reason for this, as Gause believed, is the fact, that the mathematical model does not take into account the following phenomena, that he found in his experiment. With the absence of foods, the predators did not cease to reproduce (as prescribed by the Lotka-Volterra equations system), but only, at this reproduction, the size of all new individuals becomes very small.

In our view, we can assume, all the same, that Gause fixed in his experience the one period (the last part of period) of the oscillations in number of individuals in populations, as it is prescribed by our mathematical model.

The figure 10 graphs and the graphs in figure 9 coincide qualitatively.

The repetition of the oscillations period in the Gause’s experiences has not occur, because a very small amount of bacteria involved in these experiments, and it has not allowed in experiment to display with a mathematical reliability the part of the oscillations period, when the number of persons in population (in accordance to the mathematical model) should be extremely small.

Really, the number of bacteria, that participated in Gause experience (that illustrated in figure 10) is slightly higher than the value of 1200 separate persons (in his book Gause mentions that he conducted also the experiences where the amount of bacteria reached a several thousand of individuals).

But in the presented mathematical model (figure 9) the difference in amount of persons at maximum of the population graph and at minimum of the graph (more exactly, the difference between minimal and maximal masses of population) are a six orders of magnitude. In other words, if at its minimum, the bacteria population consists of one individual, then at its maximum (for a more accurate conformity to the mathematical model) the population should consist of not less than one million persons).

It is also clear that in the practical experiences the number of bacteria can not be equal, for example, to a 10^−3^ persons; and as soon as the number of individuals is decreased to one person, the experiment ceases to conform to the mathematical model if the model prescribes at a near future the reduction in number of individuals.

Besides (if Gause would have complied in his experiments with these necessary mathematical conditions), comparing the time scale of our models, shown in figures 8 and 9, with the time scale of the figure 10, we can conclude, that for the repetition of oscillation period in Gause’s experience, he had to wait at lest a month (but the experiment, shown in figure 10, was ended after seven days of measurements).

The practicing biologists must take into account these quantitative and temporary factors mentioned above, for reliable reproduction in their own experiments these mathematical models of populations interaction.

The producing of experiments on the reproduction of predator-prey populations interaction in laboratory is associated with a greater technical difficulties that don’t allow to carry out a large number of these experiments in each laboratory, and the scientific experiences, such as this, are rare.

Amongst the experimental works of this kind the carefully executed experiences by american biologist B.G.Veilleux [4] are the most known and broadly quoted in scientific literature.

Veilleux improved the methods of conducting these experiments and has got as experiment result the stable cycles of the oscillations in the number of persons in the predator-prey populations system.

In his experiment Veilleux used as a biological model the same classical pair of bacteria: the Didinium nasutum (as predators) and the Paramecium caudatum (as prey).

Two series of these experimental data are submitted in figure 11 and 12.

**Figure 11:**
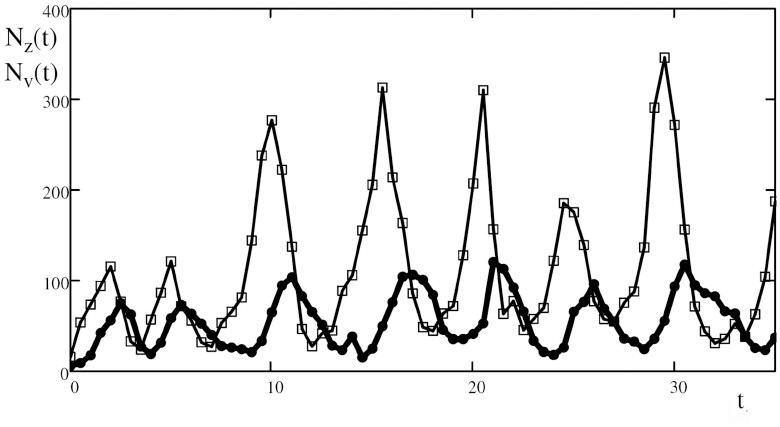
Graphs of bacteria populations interaction between predators (the Didinium nasutum bacteria) and prey (the Paramecium caudatum bacteria). The experimental data from [4] by Veilleux. Data series with the cerophyl concentration CC = 0.5 g/l.

**Figure 12:**
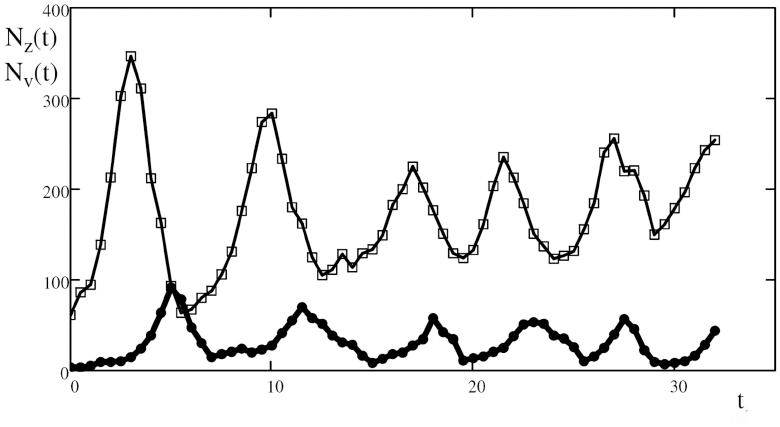
Graphs of populations interaction between predators (the Didinium nasutum bacteria) and prey (the Paramecium caudatum bacteria). The experimental data from [4] by Veilleux. Data series with the cerophyl concentration CC = 0.375 g/l.

The vertical axis represents the number of bacteria per 1 milliliter of ambience, the horizontal axis represents the time in days. The graph of predators population drawn by a thick line with the dark markers.

The Parameciums (the prey) were growing and eating in the cerophyl medium (that is the nutrient for bacteria) with different concentrations of the cerophyl.

The methylcellulose was added to a solution to obstruct the bacteria movement (for both as prey and predators). The bacteria concentration was measured twice a day, for each 12 hours, the experiments lasted 30-35 days.

The graph of predators population shown by a thick line with the dark markers, the graph of prey population shown by a hairline with the light markers.

When comparing the graphs in figures 6 and 11 we shall notice that the experimental Veilleux’s graphs repeat the main qualitative particularities of the graph 6 – the prey population has a smoothly increased amplitude of maximums of the number of individuals in population (the amplitude increases with each period); and, after the function reaches a certain limit, the general form of function remains unchanged (the form of oscillations and the period are approximately constant).

The graphs 12 and 8 also show the qualitative resemblance. The amplitude of periodic oscillations in the number of individuals decreases smoothly from the one period to the other.

This increase or reduction of oscillations amplitude – at the Veilleux’s experiments, and in our equations – corresponds to an increase or decrease in food resource for prey population.

The figures 13 and 14 show the graphs, corresponding to two theoretical models of populations interaction, that are – ceteris paribus – differ only in the amount of food resource, that is available for population of prey.

**Figure 13:**
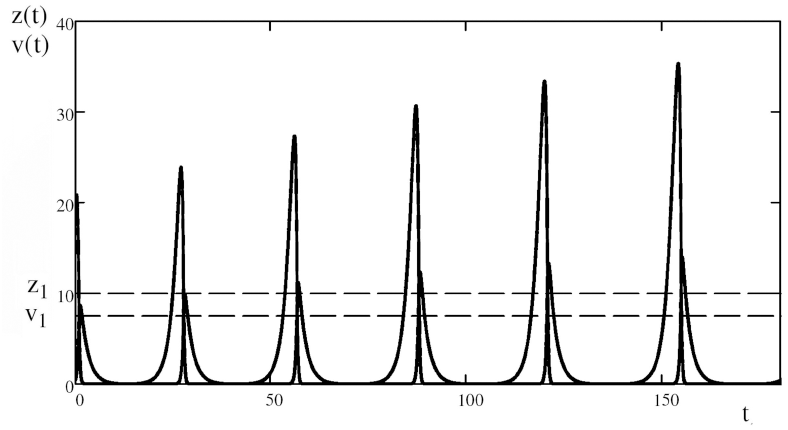
Graphs of interaction of the predators and prey populations with a food recourse quantity that is equal to a *Rz* = 2.

**Figure 14:**
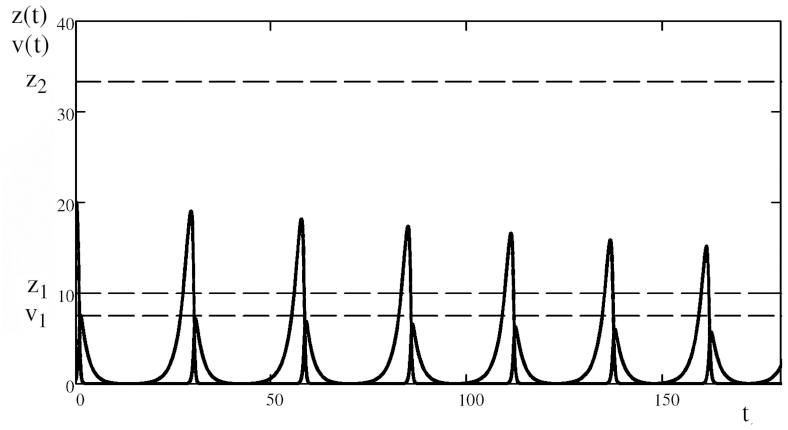
Graphs of the interaction of the predators and prey populations with a food recourse quantity *Rz* = 1.

The graphs in figure 13 are built with the following parameters values:

*k*_0*z*_ = 0.1, *k*_1*z*_ = 0.3, *k*_2*z*_ = 0.13, *k*_3*z*_ = 0.3, *k*_4*z*_ = 0.3, *k*_5*z*_ = 0.2, *k*_6*z*_ = 0.02, *α*_*z*_ = 0.8, *A*_*z*_ = 0.567, *A*_1*z*_ = 0.667, *A*_2*z*_ = 0.01, *R*_*z*_ = 2, *k*_0*v*_ = 0.1, *k*_1*v*_ = 1, *k*_3*v*_ = 0.3, *k*_4*v*_ = 0.3, *k*_5*v*_ = 0.75, *k*_6*v*_ = 0.1, *α*_*v*_ = 0.2, *A*_1*v*_ = 3, η = 0.4, *K*_1_ = 8.333, *A*_7_ = 2.5, *A*_9_ = 0.667.

The factors with the index *z* pertain to the prey population, the factors with the index *v* pertain to population of predators.

In figure 14 the value *Rz* = 2 changed to *Rz* = 1.

The value *z*_2_, which is not shown in figure 13, is *z*_2_ = 66.7.

The initial masses of the predators and prey populations are *z*_0_ = 20 and *v*_0_ = 0.5 respectively.

The numerical models of populations behaviour show that these appearing periodic oscillations of population mass can be damped noticeably.

These damped oscillations shown in figure 15, that is received with the following parameters values:

*k*_0*z*_ = 0.1, *k*_1*z*_ = 0.3, *k*_2*z*_ = 0.13, *k*_3*z*_ = 0.3, *k*_4*z*_ = 0.3, *k*_5*z*_ = 0.2, *k*_6*z*_ = 0.02, *α*_*z*_ = 6, *A*_*z*_ = 0.567, *A*_1*z*_ = 0.667, *A*_2*z*_ = 0.01, *R*_*z*_ = 2, *k*_0*v*_ = 0.1, *k*_1*v*_ = 1, *k*_3*v*_ = 0.3, *k*_4*v*_ = 0.3, *k*_5*v*_ = 0.75, *k*_6*v*_ = 0.1, *α*_*v*_ = 0.025, *A*_1*v*_ = 3, *η* = 0.4, *K*_1_ = 8.333, *A*_7_ = 2.5, *A*_9_ = 0.083.

**Figure 15:**
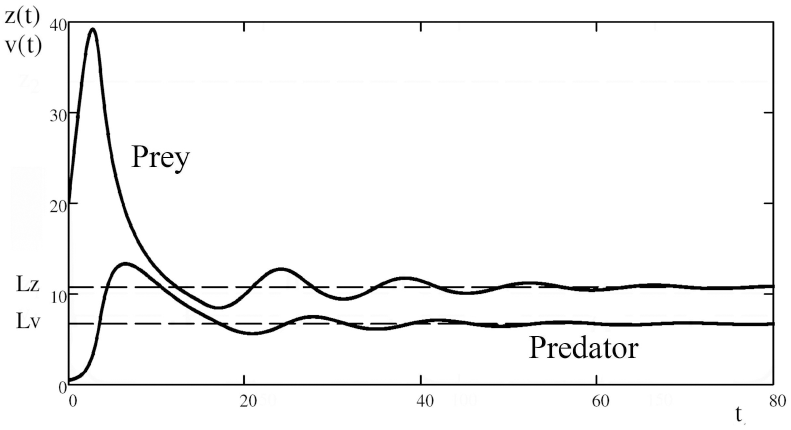
Graphs of interaction of the predators and prey populations with the damped oscillations of population mass.

The initial masses of the predators and prey populations are *z*_0_ = 20 and *v*_0_ = 0.5 respectively.

Considering that the dynamic development of the both populations ends under conditions, when the full-fed prey lives in the common territory, and the hungry predators live in the personal areas, we can use the following system of equations

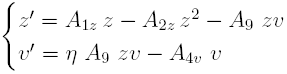

to find the development limits of population mass of the prey (*L*_*z*_) and predators (the *L*_*v*_ value).

With a replacing of left-hand sides of equations to zero, from the second equation we shall find:

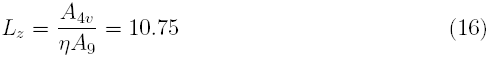

Substituting this value in the first equation, we shall obtain the value *L*_*v*_:

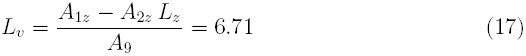

The values *L*_*z*_ and *L*_*v*_ are indicated in figure 15 by the horizontal dashed lines.

Thus, the method of mathematical simulation of the problems on the predatorprey interaction, that we have demonstrated in this article, shows us, that the laboratory experiments can be quite comparable qualitatively with the mathematical models, which, in turn, can be very useful for the practising biologist to forecast the changes in the populations development, associated with change in these or other population parameters or in the environmental conditions, both in laboratory study, and at observation of a real natural populations.

